# Mode of carbon and energy metabolism shifts lipid composition in the thermoacidophile *Acidianus*

**DOI:** 10.1101/2023.08.10.552821

**Authors:** Jeemin H. Rhim, Alice Zhou, Maximiliano J. Amenabar, Felix J. Elling, Yuki Weber, Ann Pearson, Eric S. Boyd, William D. Leavitt

## Abstract

The degree of cyclization, or ring index (RI), in archaeal glycerol dibiphytanyl glycerol tetraether (GDGT) lipids was long thought to reflect homeoviscous adaptation to temperature. However, more recent experiments show that other factors (e.g., pH, growth phase, and energy flux) can also affect membrane composition. The main objective of this study was to investigate the effect of carbon and energy metabolism on membrane cyclization. To do so we cultivated *Acidianus* sp. DS80, a metabolically flexible and thermoacidophilic archaeon, on different electron donor, acceptor and carbon source combinations (S^0^/Fe^3+^/CO_2_, H_2_/Fe^3+^/CO_2_, H_2_/S^0^/CO_2_, or H_2_/S^0^/glucose). We show that differences in energy and carbon metabolism can result in over a full unit of change in RI in the thermoacidophile *Acidianus* sp. DS80. The patterns in RI correlated with the normalized electron transfer rate between electron donor and acceptor and did not always align with thermodynamic predictions of energy yield. In light of this, we discuss other factors that may affect the kinetics of cellular energy metabolism: electron transfer chain (ETC) efficiency, location of ETC reaction components (cytoplasmic *vs*. extracellular), and the physical state of electron donors and acceptors (gas *vs*. solid). Furthermore, assimilation of a more reduced form of carbon during heterotrophy appears to decrease the demand for reducing equivalents during lipid biosynthesis, resulting in lower RI. Together, these results point to the fundamental role of the cellular energy state in dictating GDGT cyclization, with those cells experiencing greater energy limitation synthesizing more cyclized GDGTs.

**Importance:** Some archaea make unique membrane-spanning lipids with different numbers of five or six membered rings in the core structure that modulate membrane fluidity and permeability. Changes in membrane core lipid composition reflect fundamental adaptation strategies of archaea in response to stress, but multiple environmental and physiological factors may affect the needs for membrane fluidity and permeability. In this study, we tested how *Acidianus* sp. DS80 changed its core lipid composition when grown with different electron donor/acceptor pairs. We show that changes in energy and carbon metabolisms significantly affected the relative abundance of rings in the core lipids of DS80. These observations highlight the need to better constrain metabolic parameters, in addition to environmental factors, that may influence changes in membrane physiology in Archaea. Such consideration would be particularly important for studying archaeal lipids from habitats that experience frequent environmental fluctuations and/or where metabolically diverse archaea thrive.

## Introduction

The cell membrane is essential for all forms of life, serving as a physical barrier that controls the flow of nutrients and other substances to and from the external environment. Membranes also have an important bioenergetic function, as the electrochemical gradient across the cell membrane can be harnessed to conserve energy (ATP synthesis) or to perform work. Hence, the ability to modify membrane fluidity and permeability (homeoviscous adaptation) in response to changing environments is crucial for cell survival and growth. Cells can achieve this by modifying their lipid membrane composition (Chattopadhyay, 2017). While homeoviscous adaptation strategies are found across all three domains of life, Archaea provide a unique perspective to our understanding of this process. This is because archaeal lipids are fundamentally different from those of Bacteria and Eukarya, and many Archaea thrive in extreme habitats that commonly experience environmental fluctuations.

Many Archaea synthesize membrane-spanning lipids known as glycerol dibiphytanyl glycerol tetraethers (GDGTs; Figure 1). These unique structures are abundant in the core lipids produced by many thermophilic archaea (Siliakus et al., 2017) and comprise nearly all of the lipid membranes of acidophilic archaea (Macalady et al., 2004; Oger and Cario, 2013). GDGTs form very stable monolayers characterized by high packing efficiencies, high transition temperatures, and low permeabilities (Gliozzi et al., 1983; Jarrell et al., 1998; Gabriel and Chong, 2000; Nicolas, 2005). Incorporation of cyclopentane or cyclohexane rings in the alkyl cores of GDGTs further increases membrane packing and transition temperatures and helps maintain membrane integrity under extreme conditions (Gliozzi et al., 1983; Gabriel and Chong, 2000).

**Figure 1.**
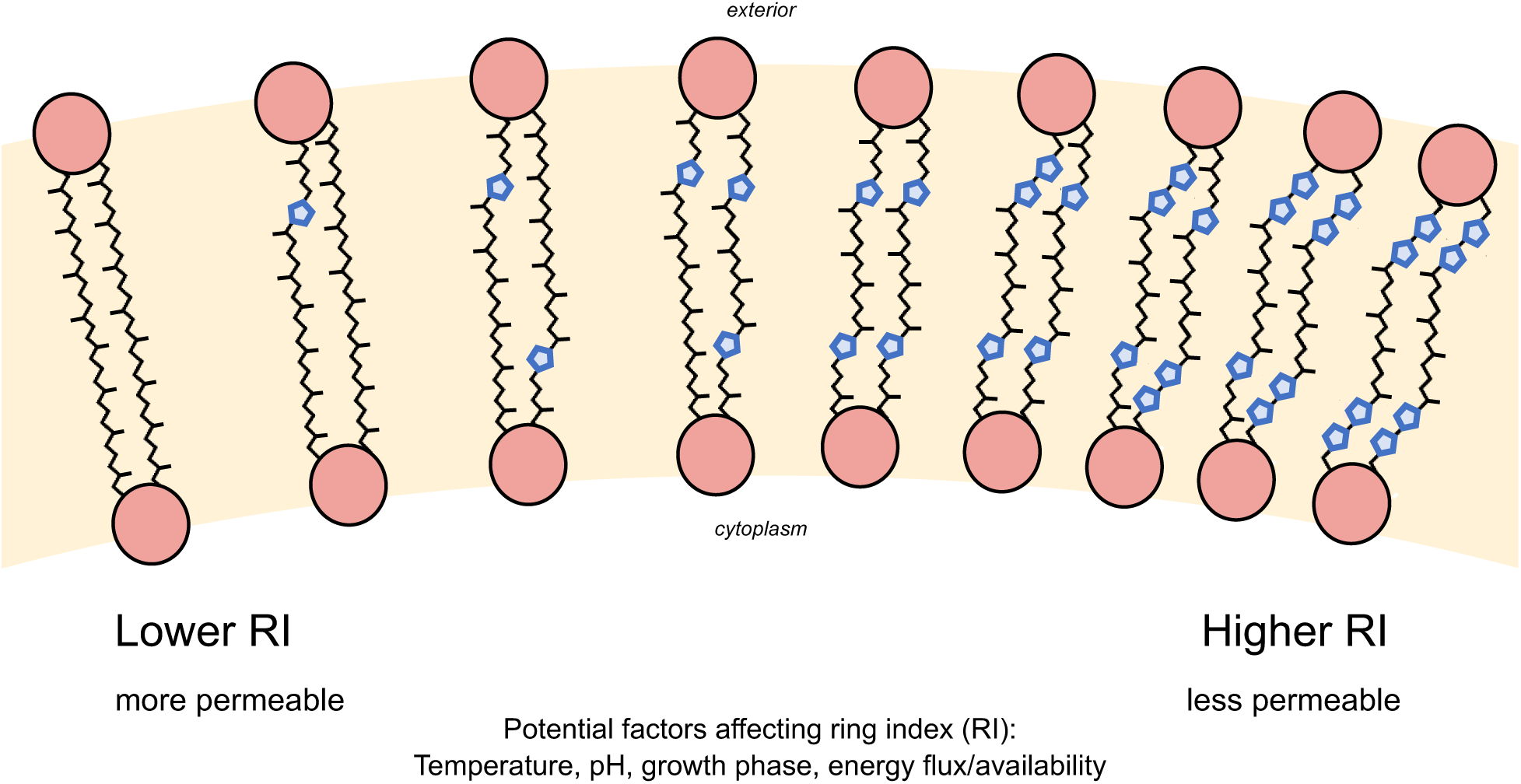
The core glycerol dibiphytanyl glycerol tetraether (GDGT) lipid structures may contain between 0 and 8 pentacyclic rings. An increase in the average number of rings per GDGT, or an increased ring index (RI), is generally associated with increased membrane packing and decreased permeability.

Besides having key physiological functions, archaeal GDGTs can serve as biomarkers that have useful applications in Earth sciences and astrobiology. Due to their high preservation potential in sediments (up to millions of years) and the information that can be inferred from the degree of cyclization, archaeal GDGTs have been widely applied as paleoenvironmental proxies (e.g., Schouten et al., 2002, 2007; Powers et al., 2010; Pearson et al., 2011). Many Archaea often belong to deeply rooted phylogenetic lineages (Pace, 1997; Schwartzman and Lineweaver, 2004; Stetter, 2006) and thus also provide contemporary analogs to study the processes that supported early life forms on Earth. Furthermore, Archaea that are well adapted to extreme environments can serve as model systems for astrobiology, as they allow us to explore the range of physicochemical conditions habitable to life (Shock and Holland, 2007).

Controlled laboratory experiments are crucial for advancing the understanding of homeoviscous adaptation in Archaea and for improving the ability to interpret GDGT biomarkers. Studies to date have investigated a variety of factors affecting the degree of cyclization in archaeal lipids. Some of these factors include temperature (De Rosa et al., 1980; Boyd et al., 2011; Schouten et al., 2013; Jain, 2014; Jensen et al., 2015; Elling et al., 2015; Feyhl-Buska et al., 2016; Cobban et al., 2020), pH (Macalady et al., 2004; Boyd et al., 2011; Elling et al., 2015; Feyhl-Buska et al., 2016; Cobban et al., 2020), ionic strength or salinity (Boyd et al., 2011; Elling et al., 2015), oxygen availability (Qin et al., 2015; Cobban et al., 2020), growth rate or energy flux (Hurley et al., 2016; Zhou et al., 2020; Quehenberger et al., 2020), and growth phase (Elling et al., 2014; Feyhl-Buska et al., 2016; Evans et al., 2018). In general, higher degrees of GDGT cyclization are associated with higher temperature, lower pH, and factors that reflect or lead to increased physiological or energetic stress (Figure 1). For example, several strains of marine ammonia-oxidizing archaea *Nitrosopumilus maritimus* and a thermoacidophile *Sulfolobus acidocaldarius* produced GDGTs with higher RIs with decreasing O_2_ availability (Qin et al., 2015; Cobban et al., 2020). RIs are also observed to increase at late growth phases in both thermoacidophilic and mesophilic archaea (Elling et al., 2014; Feyhl-Buska et al., 2016), hinting that increasing membrane packing might be a common response among Archaea to nutrient and/or energy limitation. Moreover, continuous culture experiments showed that RIs increase significantly as growth or metabolic rate and energy flux decrease (Hurley et al., 2016; Zhou et al., 2020; Quehenberger et al., 2020).

The interplay among the factors affecting membrane cyclization is particularly important for polyextremophiles that are adapted to multiple extremes, as different environmental and physiological factors could require different extents of lipid cyclization. The effects of individual parameters such as temperature, pH, ionic strength, and oxygen flux on the lipids of thermoacidophilic archaea have been investigated in previous studies (e.g., Boyd et al., 2011; Cobban et al., 2020). While these studies inform us about how RIs reflect various physicochemical factors, the effects of energy and carbon metabolism on membrane composition are poorly understood. This is partly because testing such factors requires an organism that has a flexible energy and carbon metabolism. In this study, we focus on a metabolically flexible and thermoacidophilic archaeon *Acidianus* sp. DS80 (hereafter DS80) that can use several combinations of soluble and insoluble electron donors and acceptors to support both chemolithoautotrophic and chemoheterotrophic growth (Amenabar et al., 2017, 2018; Amenabar and Boyd, 2018). DS80 was isolated from a geochemically dynamic (Colman et al., 2021) and acidic hot spring, ‘Dragon Spring’, in Yellowstone National Park, Wyoming, USA (Amenabar et al., 2017). The strain grows optimally at 80 °C and pH 3.0 (Amenabar et al., 2017). When grown autotrophically, DS80 can use molecular hydrogen (H_2_) or elemental sulfur (S^0^) as an electron donor coupled to S^0^ or ferric iron (Fe^3+^) as an electron acceptor. In addition to its ability to grow as a chemoautotroph, DS80 can also grow as an aerobic or anaerobic heterotroph. In the case of anaerobic heterotrophy, H_2_ is additionally required when S^0^ is provided as the electron acceptor (i.e., chemolithoheterotrophy; inorganic energy source with organic carbon source) (Amenabar and Boyd, 2018; Amenabar et al., 2018). The metabolic plasticity of DS80 provides a unique opportunity to investigate how different energy and carbon metabolisms in a thermoacidophilic archaeon affect lipid membrane composition.

The metabolic versatility of DS80 also raises the intriguing question as to how it selects amongst the multiple available substrates in its natural habitat to support its metabolism. Amenabar et al. (2017) characterized the thermodynamics and kinetics of substrate transformation by DS80 to explore the relationship between bioenergetics and carbon fixation efficiency. They made a counterintuitive observation where the least energetically favorable redox couple (H_2_/S^0^), or that which provided the least energy per electron transfer, resulted in the greatest degree of carbon assimilation and biomass synthesis per unit energy expended (Amenabar et al., 2017). Both of these bioenergetic parameters are likely relevant for membrane adaptations and cyclization, independent of or in concert with the established environmental drivers such as temperature and pH.

We predict that the energy available to DS80 from different electron donor/acceptor pairs and the energy demand required to access different types of electron donors and acceptors both affect core lipid composition and cyclization. To test this hypothesis, we cultivated DS80 with different electron donors and electron acceptors, both autotrophically and heterotrophically, while holding the pH of the growth medium and the incubation temperature constant. Below we document these findings with respect to studies on DS80’s energy metabolism and in light of other findings as to the role of energy availability on archaeal membrane adaptation.

## Results

*Acidianus sp.* strain DS80 was cultured as a chemolithoautotroph on three different electron donor/acceptor pairs (H_2_/S^0^, H_2_/Fe^3+^, and S^0^/Fe^3+^) with CO_2_ as the carbon source, and as a chemolithoheterotroph on H_2_/S^0^ with glucose as the carbon source (Table 1). Figure 2A shows that the rate of Fe^2+^ production decreases as the total [Fe^2+^] approaches the starting [Fe^3+^]. This suggests that Fe^3+^ was the limiting factor under Fe^3+^-reducing conditions (H_2_/Fe^3+^/CO_2_ and S^0^/Fe^3+^/CO_2_), as expected from the available electron donor and acceptor concentrations and reaction stoichiometries (Eqn. 3 and 4). On the other hand, we did not observe quantitative consumption of H_2_ under S^0^-reducing conditions (H_2_/S^0^/CO_2_ and H_2_/S^0^/glucose). For example, for the H_2_/S^0^/CO_2_ treatment, S^2–^ production decreases as the total [S^2–^] approaches 6 mM (Figure 2A); *ca*. 98 millimoles of H_2_ were initially provided in the headspace which could produce up to *ca*. 49 mM [S^2–^] upon quantitative consumption of H_2_. Thus, a factor other than electron donor/acceptor availability likely limits growth under this condition. One such factor is the production of H_2_S, which has been shown to be toxic at far lower (<80 µM) concentrations in the thermoacidophilic S^0^-reducing archaeon *Thermoproteus* strain CP80 (Urschel et al., 2016). It is possible that high concentrations of the uncharged molecule can diffuse across the membrane (Mathai et al., 2009) only to deprotonate in the more circumneutral cytoplasm (the p*K*_a_ of H_2_S/HS^−^ is 6.4 at 80 °C; Amend and Shock, 2001). This would result in acidification of the cytoplasm much like has been observed with organic acids (e.g., Aston et al., 2009).

**Figure 2.**
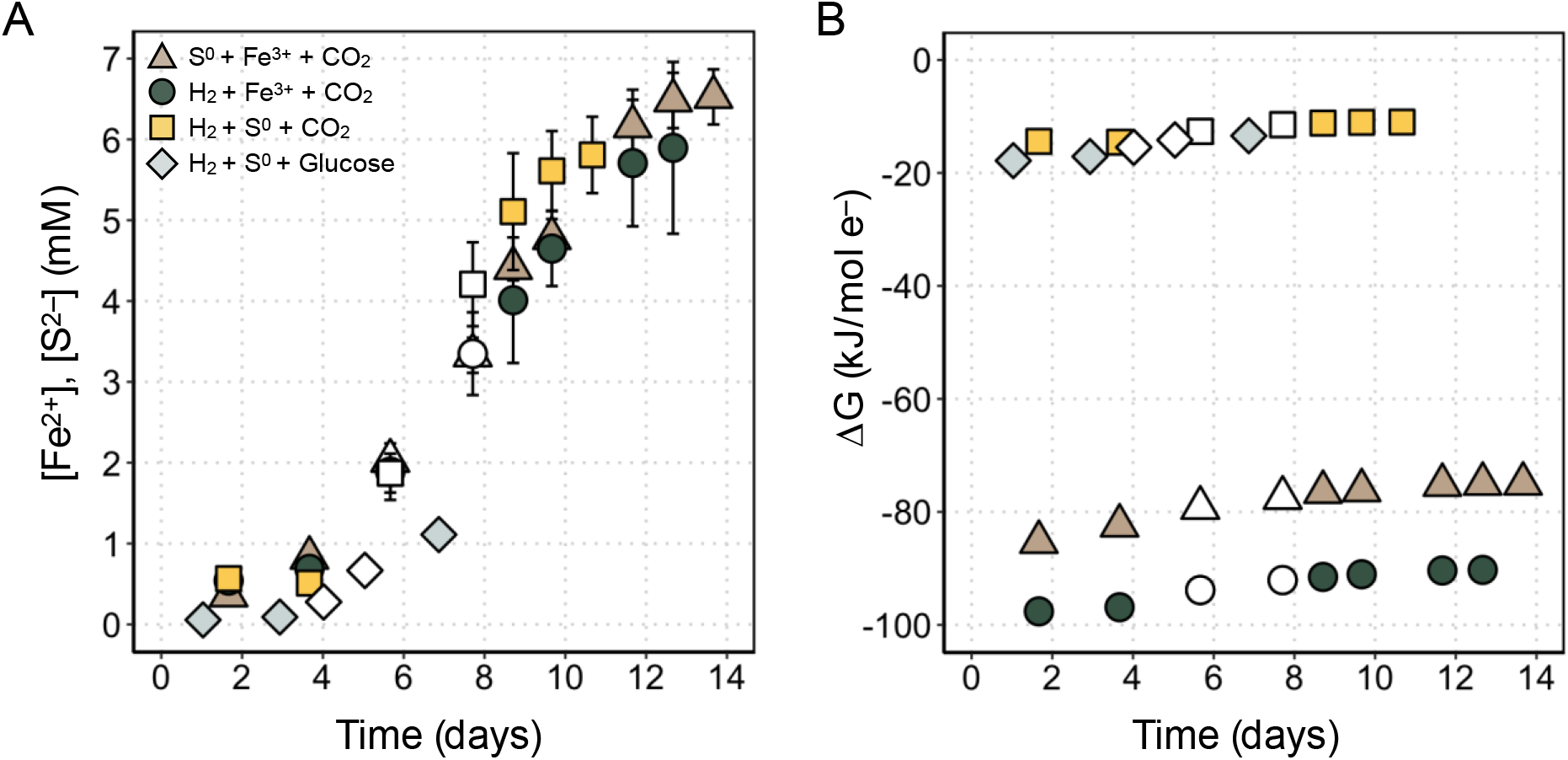
Production of metabolites and available (calculated) Gibbs free energy during growth of *Acidianus* strain DS80. (A) Changes in the concentration of metabolites (Fe_2+_ or S_2–_) as a function of incubation time. Error bars reflect the standard deviation of measurements from three biological replicates. (B) Available Gibbs free energy (ΔG) per mole of electrons transferred based on calculations and Eqn. 2–4. Open symbols represent logarithmic metabolite production (and by proxy, growth) and are the data points used to calculate ΔG_Log_ values and metabolite production rates in Table 2.

**Table 1.** Redox couples from which GDGTs were retrieved in this study.

| Experimental Condition | Electron Donor | Electron Acceptor | Carbon Source |
| --- | --- | --- | --- |
| H<sub>2</sub>/S<sup>0</sup>/CO<sub>2</sub> | H<sub>2</sub> (g) (80% v/v) | S<sup>0</sup> (s) (5 g/L) | CO<sub>2</sub> (g) (20% v/v) |
| H<sub>2</sub>/Fe<sup>3+</sup>/CO<sub>2</sub> | H<sub>2</sub> (g) (80% v/v) | Fe<sup>3+</sup> (7 mM) | CO<sub>2</sub> (g) (20% v/v) |
| S<sup>0</sup>/Fe<sup>3+</sup>/CO<sub>2</sub> | S<sup>0</sup> (s) (5 g/L) | Fe<sup>3+</sup> (7 mM) | CO<sub>2</sub> (g) (20% v/v) |
| H<sub>2</sub>/S<sup>0</sup>/glucose | H<sub>2</sub> (g) (80% v/v) | S<sup>0</sup> (s) (5 g/L) | Glucose (g) (5 mM) |

**Table 2.** Summary of bioenergetics, metabolite production rates, and ring indices across all conditions tested in this study. Available Gibbs free energy (ΔG) and rates of electron transfer and metabolite production were calculated for data points collected during logarithmic growth. Average ring indices were calculated for the biomass harvested during late logarithmic or early stationary phase. See *Supplementary Material* Table S2 for individual sample data.

| Condition | $\Delta G_{\text{Log}}^a$<br>(kJ/mol e<sup>-</sup>) | Avg. metabolite<br>production rate<br>(mmol/L/day)<sup>b</sup> | Avg. electron<br>transfer rate<br>(mmol e<sup>-</sup>/L/day)<sup>c</sup> | Inferred cell-specific<br>metabolite production rate<br>(fmol/cell/day)<sup>d</sup> | Ring Index<sup>e</sup> |
| --- | --- | --- | --- | --- | --- |
| S<sup>0</sup>/Fe<sup>3+</sup>/CO<sub>2</sub> | -78 | 0.39 ± 0.05 | 2.37 ± 0.28 | 62 ± 10 | 4.71 ± 0.03 |
| H<sub>2</sub>/Fe<sup>3+</sup>/CO<sub>2</sub> | -93 | 0.38 ± 0.08 | 0.77 ± 0.16 | 52 ± 9 | 4.10 ± 0.09 |
| H<sub>2</sub>/S<sup>0</sup>/CO<sub>2</sub> | -12 | 0.44 ± 0.16 | 0.87 ± 0.32 | 7 ± 1 | 3.89 ± 0.14 |
| H<sub>2</sub>/S<sup>0</sup>/glucose | -15 | 0.10 ± 0.04 | 0.20 ± 0.09 | n/a | 3.61 ± 0.11 |

<sup>a</sup> Average $\Delta G$ values for data points representing logarithmic increase in metabolite concentration (open symbols in Figure 2). For the H<sub>2</sub>/S<sup>0</sup>/glucose condition, it is assumed that glucose does not significantly contribute to energetic $\Delta G$ . See 2.4 for a detailed description of values used for $\Delta G$ calculations.
<sup>b</sup> Rates of Fe<sup>2+</sup> production per liter for the S<sup>0</sup>/Fe<sup>3+</sup>/CO<sub>2</sub> and H<sub>2</sub>/Fe<sup>3+</sup>/CO<sub>2</sub> conditions, based on the Ferrozine assay results; rate of S<sup>2-</sup> production per liter for the H<sub>2</sub>/S<sup>0</sup>/CO<sub>2</sub> condition, based on the methylene blue assay results (dissolved concentration) and applying a standard gas-phase equilibrium correction to calculate total sulfide.
<sup>c</sup> Rates of electrons transferred per liter of culture, based on metabolite production rates and reaction stoichiometries in Eqn. 2–4.
<sup>d</sup> Metabolite production rates were divided by inferred cell density values. Cell density values were calculated from measured metabolite concentrations, [Fe<sup>2+</sup>] or [S<sup>2-</sup>], and the growth yield calculated for logarithmic growth from data reported in Amenabar et al. (2017): $2.5 \times 10^{12}$ cells per mole of Fe<sup>2+</sup> produced for S<sup>0</sup>/Fe<sup>3+</sup>/CO<sub>2</sub>; $3.0 \times 10^{12}$ cells per mole of Fe<sup>2+</sup> produced for H<sub>2</sub>/Fe<sup>3+</sup>/CO<sub>2</sub>; $20.6 \times 10^{12}$ cells per mole of S<sup>2-</sup> produced for H<sub>2</sub>/S<sup>0</sup>/CO<sub>2</sub>.
<sup>e</sup> Ring indices were calculated from biomass harvested during late logarithmic or early stationary phase.

The lipid compositions of DS80 vary across different growth conditions. The degree of cyclization (i.e., RI) changed consistently with metabolite production rates and electron transfer rates (Table 2; Figure 3). Overall, the core lipid profiles of DS80 primarily consisted of GDGT-4 and GDGT-5. The Fe^3+^-reducing conditions (H_2_/Fe^3+^/CO_2_ and S^0^/Fe^3+^/CO_2_) resulted in higher relative abundances of GDGT-4, GDGT-5, and GDGT-6 than the S^0^-reducing conditions (Figure 3). Consequently, Fe^3+^-reducing conditions resulted in higher RI values compared to S^0^-reducing conditions (H_2_/S^0^/glucose and H_2_/S^0^/CO_2_). There was more than a full unit difference between the highest (4.71 ± 0.01) and lowest (3.61 ± 0.08) RIs observed in cultures grown on S^0^/Fe^3+^/CO_2_ and H_2_/S^0^/glucose, respectively (Table 2; Figure 3). Between the two S^0^-reducing conditions, the heterotrophic condition resulted in a significantly lower RI (3.61 ± 0.08; H_2_/S^0^/glucose) compared to the autotrophic condition (3.89 ± 0.14; H_2_/S^0^/CO_2_), although the calculated available Gibbs free energy during log phase (termed ΔG_Log_ herein) for the two conditions were comparable. Overall, the trends observed in RIs were better explained by the normalized electron transfer rate (R^2^ = 0.94; Figure 4B) than by the ΔG_Log_ values (R^2^ = 0.45; Figure 4A).

**Figure 3.**
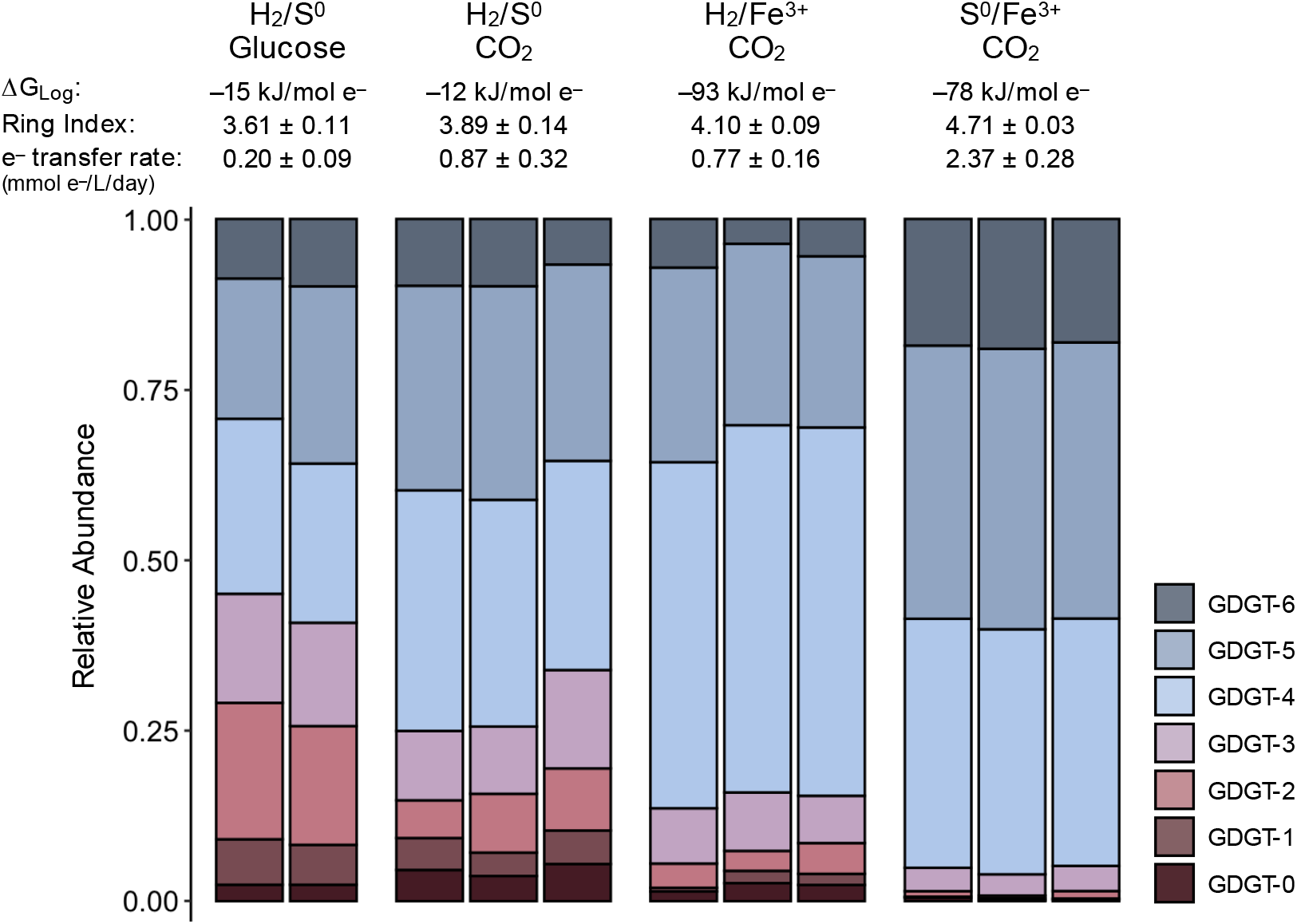
Relative abundances of core GDGT lipids as a function of electron donor/electron acceptor couples and carbon sources provided. The available Gibbs free energy during log phase (ΔG_Log_), average ring index (the weighted average degree of cyclization), and normalized electron transfer rates for each condition are shown above corresponding stacked bar charts.

**Figure 4.**
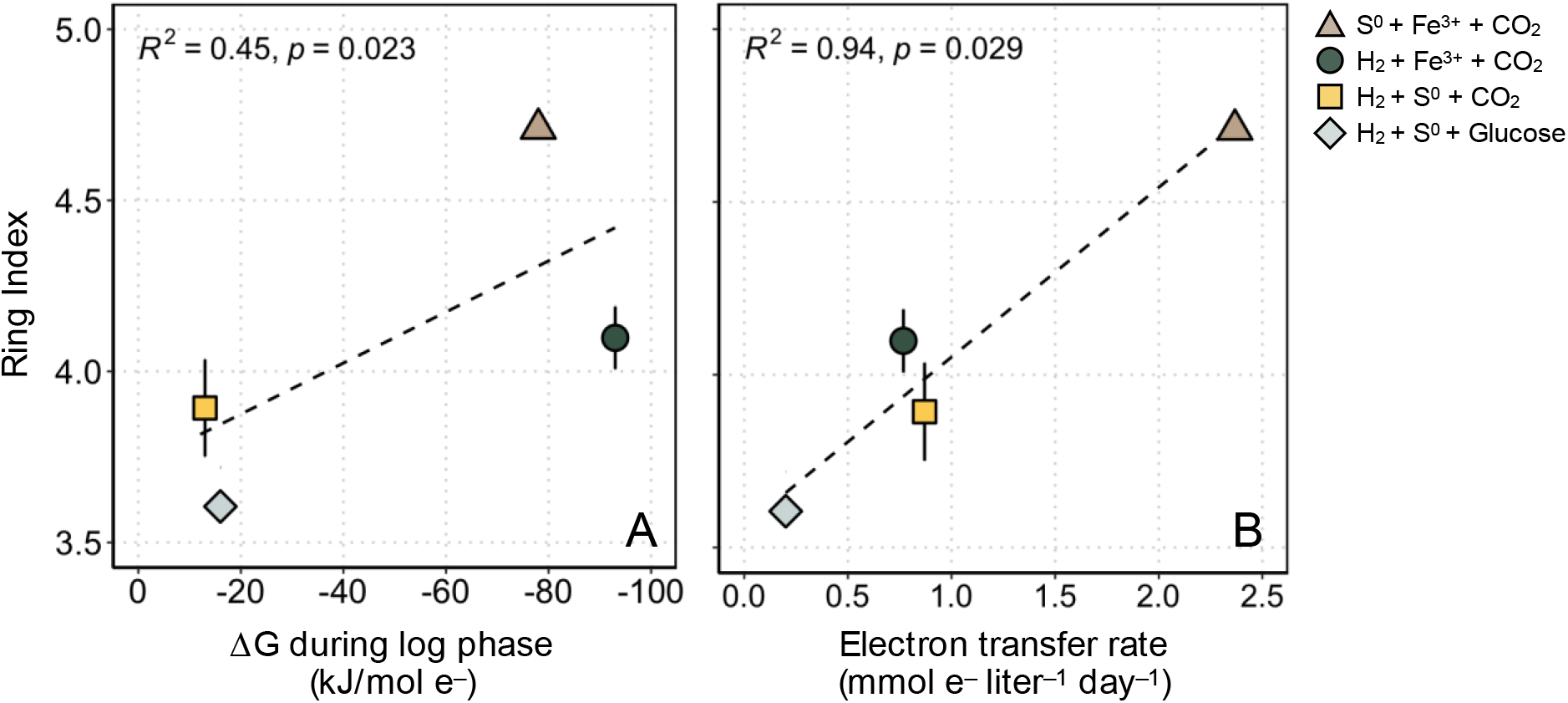
Average ring index values as a function of available Gibbs free energy and electron transfer rate. (A) The ΔG values reflect calculations made during logarithmic growth (ΔG_Log_) for each condition tested in this study (see 2.4). (B) Electron transfer rates were calculated from the measured rates of Fe_2+_ or S_2–_ production per liter of culture during logarithmic growth and using reaction stoichiometries in Eqn. 2–4. The R_2_ value in each panel is for the regression line (dashed line) for all average ring index values.

## Discussion

For (poly)extremophilic archaea (Capece et al., 2013), the balance between bioenergetics and membrane plasticity becomes particularly important for survival and growth. On one hand, ring synthesis reduces the demand for electrons (and thus electron donors/acceptors), as the reduction of double bonds during saturation (opposite of cyclization) requires a net expenditure of reducing equivalents (Hurley et al., 2016; Pearson, 2019). As such, under electron donor or acceptor (energy or electron) limitation, a higher degree of cyclization helps to minimize the expenditure of reducing equivalents during lipid synthesis and provides tighter membrane packing that reduces leakage of ions across the cell membrane (Hulbert and Else, 2005). On the other hand, factors such as the nature of electron donors or acceptors (e.g., soluble *vs*. insoluble or charged *vs*. uncharged) may impose an opposing biosynthetic pressure to maintain a lower degree of cyclization. A recent study reported changes in bacterial lipid composition as a function of electron acceptor (nitrate *vs*. manganese dioxide), and the changes were attributed to the differences in the electron transport chains (ETC) embedded in the cell membrane (Ding et al., 2022). For archaeal lipids, changes in the membrane composition in response to the extent of anaerobiosis (e.g., S^0^ *vs*. O_2_ as the electron acceptor) have been observed (Trincone et al., 1989). While homeoviscous adaptations in Archaea have been studied in the context of different environmental and physiological forcings (De Rosa et al., 1980; Boyd et al., 2011; Schouten et al., 2013; Jain, 2014; Elling et al., 2014, 2015; Jensen et al., 2015; Qin et al., 2015; Hurley et al., 2016; Feyhl-Buska et al., 2016; Zhou et al., 2020; Quehenberger et al., 2020), it remains poorly understood how cells accommodate bioenergetic needs while balancing needs for different electron donors/acceptors and how this is reflected in cyclization patterns in GDGT lipids.

In this study, the metabolic versatility of DS80 allowed us to begin to investigate archaeal membrane GDGT cyclization patterns during growth on various electron donor/acceptor pairs while keeping temperature, pH, and growth phase at the time of biomass collection consistent across treatments. When the thermodynamic predictions of energy yield from the tested redox couples are combined with the bioenergetics argument to increase membrane packing (i.e., increased RI) under energy limitation, we expect the highest RI to be measured in culture conditions with the least favorable ΔG_Log_ values (H_2_/S^0^/CO_2_ and H_2_/S^0^/glucose). Counterintuitively, we observed that the H_2_/S^0^/CO_2_ and H_2_/S^0^/glucose treatments yielded the lowest RIs averaging 3.89 ± 0.14 and 3.61 ± 0.08, respectively (Table 2; Figure 3). Interestingly, the treatment with the most favorable ΔG_Log_ value (H_2_/Fe^3+^/CO_2_) yielded intermediate RIs averaging 4.10 ± 0.09, ranking after only the S^0^/Fe^3+^/CO_2_ treatment with RIs averaging 4.71 ± 0.03 (Table 2; Figure 3). The difference of over a full unit of RI observed in this study is comparable to the magnitude of changes observed upon significant shifts in temperature (15 to 20 °C shift) or pH (e.g., nearly 2 log shift) (Shimada et al., 2008; Boyd et al., 2011).

We propose that the trends in membrane cyclization patterns in DS80 reflect differences in energy demand associated with the nature and availability of electron donors and acceptors that are not accounted for in thermodynamic calculations. In general, we observed comparatively lower RIs during S^0^ reduction compared to those during Fe^3+^ reduction. In the case of carbon assimilation and biomass production, Amenabar et al. (2017) identified the difference in electron transfer efficiency as the primary factor underlying the counterintuitive observation of the highest biomass yield during S^0^ reduction. For the H_2_/S^0^ redox couple, electrons are transferred via an extremely short ETC involving a membrane-bound [NiFe]-hydrogenase and sulfur reductase (SRE) complex linked by a quinone cycle (Laska et al., 2003). Interestingly, electron transfer efficiency was significantly correlated with membrane cyclization in DS80 as determined herein, where lower electron transfer efficiency during Fe^3+^ reduction is compensated by higher cyclization to presumably decrease the expenditure of reducing equivalents when compared to growth on S^0^. This suggests that the nature of electron donors and acceptors, and microbial adaptations to extract or donate electrons to those substrates, both affect lipid composition.

The treatments with the lowest and highest RIs (H_2_/S^0^/glucose and S^0^/Fe^3+^/CO_2_, respectively; Table 2; Figure 3) can be further examined in the context of the location of where the redox reactions (electron transfers) are taking place. The membrane-bound SRE complex that facilitates sulfur reduction in a closely related strain *Acidianus ambivalens* (and likely DS80) is extracellularly-oriented, meaning that reduction takes place outside of the cell (Laska et al., 2003; Amenabar et al., 2018; Counts et al., 2021). In contrast, the unique sulfur oxygenase reductase (SOR) complex facilitates cytoplasmic oxidation of sulfur in a few lineages of *Sulfolobales* including *Acidianus* (Ulrich et al., 2006; Kletzin, 2008; Counts et al., 2021). The protein responsible for S^0^ transport into the cell remains unidentified, and the source of intracellular sulfur for the SOR system is not well understood. It has been suggested that the sulfide:quinone oxidoreductase (SQO) may serve as the intracellular sulfur source by forming polysulfide chains from the oxidation of hydrogen sulfide (Counts et al., 2021). Whether it is via an unknown transporter for S^0^ and/or via polysulfide formation by SQO, cytoplasmic sulfur oxidation via the SOR system would require additional energy expenditure (i.e., higher RI is favorable) to account for sulfur acquisition. Further, the number of enzymes/proteins involved in oxidation of sulfur via SOR to generate energy (24 subunits; Kletzin, 2008) versus reduction of sulfur via SRE (5 subunits; Laska et al., 2003) would impose an additional biosynthetic burden on cells oxidizing S^0^. Besides the SOR complex, many other enzymes are required during the oxidative metabolism of S^0^ that would further increase energy demands associated with extracting electrons from this substrate. Overall, the energy expenditures expected for intracellular sulfur oxidation and extracellular sulfur reduction align with the highest and lowest RIs observed from the S^0^/Fe^3+^/CO_2_ and H_2_/S^0^/glucose conditions, respectively.

In addition to the additional energy expenditures associated with S^0^ oxidation, lower energy efficiency during Fe^3+^ reduction may also contribute to the high RI observed in the S^0^/Fe^3+^/CO_2_ treatment. Previous work has shown that DS80 can reduce soluble Fe^3+^ produced by acidic dissolution of ferric iron minerals (Amenabar and Boyd, 2018) or directly reduce Fe^3+^ on the mineral surface (Chanda et al., 2021) during dissimilatory iron reduction (DIR). Electron transfer likely takes place outside of the cell during DIR given that DS80 produces pili-like structures exclusively during DIR and that it lacks biochemically characterized or bioinformatically identified genes encoding for *c*-type cytochromes (Amenabar et al., 2017). Moreover, intracellular production of Fe^2+^ during DIR would likely be toxic to cells (Bennett and Gralnick, 2019). While the exact location and mechanism of the electron transfer to Fe^3+^ in DS80 remains to be determined, extracellular electron transfer during DIR may be possible through a non-dedicated mechanism where electrons diverted from the membrane spontaneously reduce Fe^3+^. A similar mechanism has been shown in methanogens during pyrite (FeS_2_) reduction, where membrane electrons with low (< 280 mV) redox potentials are diverted toward the reduction of FeS_2_ (Spietz et al., 2022). Regardless of the exact mechanism, these plausible routes of extracellular electron transfer during DIR are likely to be inefficient and may further contribute to energy limitation that leads to the highest RI observed in the S^0^/Fe^3+^/CO_2_ treatment.

Another consideration for the treatments with the lower RIs (H_2_/S^0^/glucose and H_2_/S^0^/CO_2_) is the electron donor acquisition process for molecular hydrogen (gas). The SRE complex for sulfur reduction is accompanied by a [NiFe]-hydrogenase that catalyzes the interconversion of H_2_ to protons and electrons (Ogata et al., 2016). A full suite of genes encoding a membrane-bound [NiFe]-hydrogenase (HynSL) has been identified in the genome of DS80 (Amenabar et al., 2018). Although the location of this [NiFe]-hydrogenase in DS80 is not well understood, most known membrane-bound hydrogenases are facing the cytoplasmic side of the membrane (Leul et al., 2005). When this is the case, enzyme activity is reliant on diffusion of dissolved H_2_ across the cell membrane and into the active site of the hydrogenase. This passive diffusion step may set upper bounds on RI and membrane compactness, as tightly packed membranes with high RIs could restrict the transmembrane diffusion of H_2_. In this respect, the need to maintain membrane fluidity during H_2_ oxidation may place an additional biosynthetic pressure conducive to lower RIs for the H_2_/S^0^/glucose and H_2_/S^0^/CO_2_ treatments.

Besides the impact of electron donors/acceptors and associated electron transfer pathways, carbon metabolism also appears to contribute to changes in membrane cyclization. It has been shown that, when S^0^ is provided to DS80 as the electron acceptor, inorganic and organic carbon sources could only support growth when H_2_ was also provided (Amenabar et al., 2018). Considering the obligate requirement for H_2_ during chemolithoheterotrophic metabolism during S^0^ reduction and the insignificant difference in the yield of cells when grown with H_2_/CO_2_/S^0^ and compared to H_2_/CO_2_/S^0^ supplemented with acetate (Amenabar et al., 2018), we assumed no significant energetic ΔG contribution from glucose for the H_2_/S^0^/glucose treatment. Accordingly, the ΔG_Log_ values estimated for the H_2_/S^0^/CO_2_ and H_2_/S^0^/glucose conditions are comparable (Figure 2B; Table 2) and thus the RIs would be expected to be comparable if they were only influenced by this parameter. However, it is possible that some or all of the carbon used for lipid biosynthesis originates from glucose during chemolithoheterotrophic growth. When lipid synthesis starts from a more reduced form of carbon, the demand for reducing equivalents during lipid synthesis would be lower. This is in line with the difference in RIs observed between the autotrophic H_2_/S^0^/CO_2_ (3.89 ± 0.14) and heterotrophic H_2_/S^0^/glucose (3.61 ± 0.08) conditions (Table 2; Figure 3).

Overall, the additional considerations above provide a better explanation for the cyclization patterns observed in this study (Figure 3). These collective considerations are also consistent with the observation that RIs correlate better with electron transfer rates (R^2^ = 0.94) than with calculated ΔG_Log_ values (R^2^ = 0.45) (Figure 4), as metabolic rates reflect a net physiological response to environmental forcings. While cell abundance was not quantified in this study, the growth yield for each autotrophic condition can be inferred from the data reported in Amenabar et al. (2017). Assuming the same growth yield for the autotrophic and heterotrophic H_2_/S^0^ treatments (H_2_/S^0^/CO_2_ and H_2_/S^0^/glucose), the correlation is still significant (R^2^ = 0.94) between RIs and cell-specific electron transfer rates (i.e., metabolic rates normalized to both reaction stoichiometry and biomass yield) (Supplementary Material, Figure S2).

The lower RIs at lower electron transfer rates observed in this study (Figure 4B) appear to contradict observations from previous studies where higher RIs were observed at lower metabolic rates or stationary growth phase. However, upon closer consideration, the combined observations reveal a broader link between energy limitation and membrane cyclization. In experiments with the thermoacidophile *S. acidocaldarius*, both in fed-batch cultures with electron acceptor limitation and continuous cultures with electron donor limitation, slower cellular doubling times yielded GDGTs with higher RIs (Zhou et al., 2020; Quehenberger et al., 2020; Cobban et al., 2020). Similarly, in continuous cultures of *N. maritimus* with electron donor limitation, slower growth and cell-specific ammonia oxidation rates resulted in higher RIs (Hurley et al., 2016). Both *S. acidocaldarius* and *N. maritimus* continuous culture experiment results are consistent with previous batch culture experiments, where lag or stationary growth phase (i.e., proxy for lower metabolic rates) resulted in higher RIs (Elling et al., 2014; Feyhl-Buska et al., 2016; Evans et al., 2018). The general trends from prior batch and continuous culture studies are directly in line with the bioenergetics prediction of increased membrane packing (higher RI) under energy limitation (slower metabolic rate or lag/stationary growth phase). Unlike other batch experiments, biomass was harvested at about the same growth phase across experiments in this study. Furthermore, DS80 was grown on a range of different electron donor/acceptor pairs to test the effect of energy and carbon metabolism on membrane cyclization, whereas the batch and continuous culture studies noted above tested the effect of metabolic rate or growth phase within the same metabolic regime. The trend we observe reflects the differential energy limitation experienced by DS80 in the broader context of differences — in ETC efficiency, location of redox chemistry, and physical state of substrate — across energy and carbon metabolism modes. Together, the work here with DS80 and prior studies across GDGT-producing archaea highlight the role of carbon and energy metabolism (electron transfer efficiency, in particular) play in shaping GDGT cyclization and archaeal membrane physiology in nature.

In this study, we used a metabolically flexible thermoacidophilic archaeon *Acidianus* sp. DS80 to test the effects of energy and carbon metabolism on membrane cyclization. Experimental results revealed that both the metabolite production rate and the degree of cyclization varied across growth treatments. However, the patterns in RI did not always correlate with the ΔG_Log_ values for corresponding redox couples. We discussed several factors that may affect reaction kinetics during the growth of DS80 and that may influence RI. The generally lower RI observed in S^0^-reducing conditions can be attributed to the: 1) high efficiency of the short electron transfer pathway involving a [NiFe]-hydrogenase and sulfur reductase linked by quinone, 2) favorable location of redox chemistry during S^0^ reduction, and 3) potential need to maintain membrane fluidity for substrate diffusion during H_2_ oxidation. Moreover, carbon metabolism appears to affect membrane cyclization, where assimilation of a more reduced form of carbon (glucose *vs*. CO_2_) during heterotrophy results in lower RI compared to autotrophic conditions. Altogether, these findings highlight the effects of energy and carbon metabolism on membrane cyclization in DS80 in the same or opposite direction of the biosynthetic pressure imposed by extreme temperature and pH. Taken together, factors that increase the energetic demands on cells (less efficient ETCs, autotrophic metabolism, unfavorable locations of proteins involved in redox chemistry, suboptimal pH and temperature) appear to generally result in increased cyclization of GDGT lipids (i.e., GDGT RI). Further understanding the complex interplay among environmental and physiological factors that influence patterns of GDGT cyclization will improve the application of archaeal GDGTs as records of past environments.

## Materials and Methods

### Strain and cultivation procedures

To examine the effect of carbon sources, electron donors, and electron acceptors on the composition of GDGT lipids, axenic cultures of DS80 were grown on defined mineral medium (Boyd et al., 2007) consisting of NH_4_Cl (0.33 g/L), KCl (0.33 g/L), CaCl_2_ • 2H_2_O (0.33 g/L), MgCl_2_ • 6H_2_O (0.33 g/L), and KH_2_PO_4_ (0.33 g/L). Following autoclave sterilization, filter-sterilized Wolfe’s vitamins (1 mL/L) and SL-10 trace metals (1 mL/L) were added to the base mineral medium. Electron donors and electron acceptors were then added after autoclave sterilization, according to each experimental condition (Table 1). Elemental sulfur (S^0^; sulfur precipitated powder, EMD Millipore) was sterilized by baking at 100 °C for 24 hours and added to the medium at a concentration of 5.0 g/L. Ferric iron (Fe^3+^) was added in the form of ferric sulfate solution to a final concentration of 7 mM. Importantly, DS80 is unable to respire sulfate but can respire ferric iron; sulfate from ferric sulfate or sulfide (from S^0^ reduction) can serve as sulfur sources for DS80 (Amenabar et al., 2017).

The overall medium preparation for autotrophic cultures (H_2_/S^0^, H_2_/Fe^3+^, and S^0^/Fe^3+^) followed a previously described protocol involving 2 hours of purging with sterile N_2_ gas followed by replacement of the headspace with sterile H_2_/CO_2_ (80:20, v/v) or H_2_/N_2_ (80:20, v/v) (Amenabar et al., 2017, 2018). Three biological replicates were prepared for each experimental condition in 5-liter glass bottles (Fisherbrand, FB-800-5000), each with a final liquid volume of 2 liters and sealed with butyl rubber stoppers. Each bottle was inoculated with 200 mL of a log phase culture of DS80 and incubated statically at 80 °C (Binder Avantgarde BD56). The pH of the growth medium was set at an initial value of 3.0 for all conditions and remained within 0.1 units throughout the experiment (data not shown). Medium preparation for heterotrophic cultures (H_2_/S^0^/glucose) followed the general procedure described above and was distributed into smaller individual 160-mL serum bottles, each with a final liquid volume of 100 mL. Each serum bottle was additionally amended with sterile glucose solution to a final concentration of 5 mM.

Based on the stoichiometry shown in Eqn. 2a, 5 g/L of S^0^ would meet the theoretical requirement for complete oxidation of the H_2_ supplemented in S^0^-reducing conditions (H_2_/S^0^/CO_2_ and H_2_/S^0^/glucose). Based on the stoichiometry shown in Eqn. 3 and 4, 5 g per L of S^0^ and 80% H_2_ in the 2.88 L headspace would meet the theoretical requirement for complete reduction of the Fe^3+^ supplemented in Fe^3+^-reducing conditions (S^0^/Fe^3+^/CO_2_ and H_2_/Fe^3+^/CO_2_).

### Measurement of metabolic activities

Growth of microbial cultures is traditionally assessed using direct cell counts or optical density measurements. In this study, the production of total sulfide or ferrous iron was used as a proxy for microbial growth since they have previously been shown to correlate strongly and positively with cell densities (Amenabar et al., 2017, 2018). Concentrations of dissolved sulfide were determined via the methylene blue reduction method (Fogo and Popowsky, 1949) for H_2_/S^0^ cultures. The amount of total sulfide produced was calculated from the dissolved concentrations using standard gas-phase equilibrium (Eqn. 2b) calculation (Boyd et al., 2007). Concentrations of reduced iron (Fe^2+^) were determined via the ferrozine assay (Viollier et al., 2000) for cells provided with H_2_/Fe^3+^ or S^0^/Fe^3+^ (*Supplementary Material*, Figure S1). Metabolic rates were calculated based on the results of the aforementioned assays (*Supplementary Material*, Figure S1).

### Lipid analyses

All DS80 biomass samples were harvested via filtration upon reaching late logarithmic or early stationary phase, based on measurements of metabolites and comparisons to previous growth curves (Amenabar et al., 2017, 2018). Cultures were removed from the incubator and rapidly cooled to room temperature in 4 °C water baths. Cooled samples were then filtered onto 0.22 μm pore size glass fiber filters (Advantec GF7547MM Grade GF75 Glass Fiber Filters, GFF; 47 mm diameter). Prior to use, all glass components were combusted at 350 °C for 4 h. To remove the bulk of solid sulfur and/or iron (oxy)hydroxide precipitates, cultures were decanted into 250 mL centrifuge bottles and spun down gently (3 min at 600 *g*). The resulting supernatant containing suspended cells was concentrated onto glass fiber filters and frozen at –80 °C until lipid extractions.

Prior to extraction, filters were cut into small pieces using pre-combusted stainless-steel scissors. 100 ng of synthetic C_46_ GDGT standard (Patwardhan and Thompson, 1999) was added to each sample for quantification. Samples were hydrolyzed in 5% (v/v) methanolic HCl (70 °C, 90 minutes) to convert intact polar lipids to core lipids. Following acid hydrolysis, core lipids were extracted by ultrasonication after the addition of either dichloromethane (DCM, used for H_2_/S^0^, H_2_/Fe^3+^, and S^0^/Fe^3+^ samples) or methyl tert-butyl ether (MTBE, used for H_2_/S^0^/glucose samples). Phase separation was induced with either a 1:1 mixture of DCM and water (H_2_/S^0^, H_2_/Fe^3+^, and S^0^/Fe^3+^ samples) or with hexane (H_2_/S^0^/glucose samples). Core GDGTs were then purified over activated aluminum oxide by elution with DCM/methanol (1:1, v/v). The resulting fraction was dried under a flow of N_2_, resuspended in 500 μL hexane/isopropanol (99:1, v/v), passed through a 0.45-μm PTFE filter, and stored at –20 °C until analysis.

The extracted core GDGTs were analyzed by ultra high performance liquid chromatography-atmospheric pressure chemical ionization-mass spectrometry (UHPLC-APCI-MS) using an Agilent 1290 Infinity series UHPLC system coupled to an Agilent 6410 triple-quadrupole MS, operated in positive mode (gas temperature: 350 °C; vaporizer temperature: 300 °C; gas flow: 6 L min^-1^, nebulizer pressure: 60 psi). Core lipids in the filtered extract were separated using normal phase liquid chromatography-mass spectrometry. Analytical separation of GDGTs was achieved by injecting 10 μL of total lipid extract onto a Prevail Cyano column maintained at 50 °C. GDGTs were eluted using a linear gradient from 0.2% to 10% (v/v) isopropyl alcohol (IPA) in hexane at a flow rate of 0.5 mL/min as previously described (Becker et al., 2013). At the end of each sample run, the columns were back-flushed with a 70:30 mixture of hexane:IPA (90:10, v/v) and IPA:methanol (70:30, v/v). Columns were re-equilibrated to initial conditions before proceeding with the next sample run. The MS was operated in single ion monitoring mode (dwell time 25 ms, fragmentor voltage: 75 V) and GDGTs were quantified by integration of the ion chromatograms of analytes relative to the C_46_ internal standard peak. Peak areas of all GDGT species are provided in *Supplementary Material*, Table S1.

Ring index (RI) was calculated for each sample according to the formula, following (Pearson et al., 2004). The definition below accounts for GDGTs with up to 6 rings (GDGT-0 to GDGT-6), as GDGT-7 and GDGT-8 were not detected:

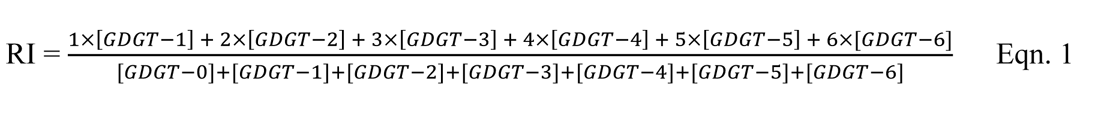

Note that these results do not consider the relative contribution of calditol-linked GDGTs. Zeng et al. (2018) observed that calditol-linked lipids are slightly more cyclized compared to GDGTs in *S. acidocaldarius*. The acid hydrolysis method used in this study does not remove the ether-bound calditol head group, thus the RIs reported in this study may be underestimated. Based on previous observations (e.g., Zeng et al., 2018; Quehenberger et al., 2020; Cobban et al., 2020), we assume that the relative offset in RIs between experimental conditions, especially among autotrophic conditions, likely will remain consistent between the calditol-linked GDGT pool and the remaining GDGT pool (Cobban et al., 2020).

### Bioenergetic calculations

Each experimental growth condition was dependent on one of the following chemical reactions in medium at pH 3.0:

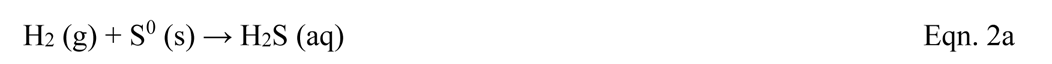

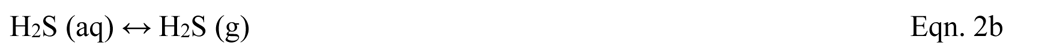

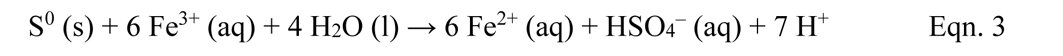

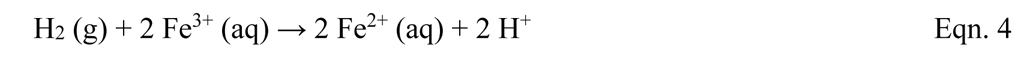

The amount of free energy available during each of these chemical reactions was calculated using the following equation to account for non-standard conditions:

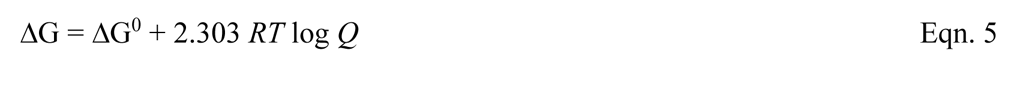

where ΔG is the Gibbs free energy of reaction (J mol^−1^); ΔG^0^ is the standard Gibbs free energy of reaction (J mol^−1^); *R* is the ideal gas constant (8.314 J mol^−1^ K^−1^); *T* is the temperature in K; and *Q* is the reaction quotient. Because these are dilute solutions, activity coefficients were assumed to be 1 for all dissolved compounds (Spear et al., 2005). [S^2–^] and [Fe^2+^] values were calculated based on spectrophotometric measurements, and [Fe^3+^] values were calculated from the starting [Fe^3+^] and measured [Fe^2+^] values. We did not quantify dissolved H_2_ concentrations and assumed a constant [H_2_] in equilibrium with *ca*. 1 atmosphere of *p*H_2_ (*ca*. 768 µM). The values reported in Table 2 were calculated by taking the average between data points from two time points representing the logarithmic growth (open symbols in Figure 2).

## Acknowledgements

Financial support was provided by ACS-PRF DNI grant #57209-DNI2 (WDL, YW), the Walter & Constance Burke Fund at Dartmouth College (WDL), the NASA NH Space grant NNX15AH79 (WDL, AZ), NSF EAR 1928303 (WDL), the Gordon and Betty Moore Foundation and NSF-1843285 (AP), NSF EAR 1820658 (EB), the Deutsche Forschungsgemeinschaft grant 441217575 (FJE), and the Dartmouth Society of Fellows (JHR). We thank the other members of the Leavitt, Boyd and Pearson labs for thoughtful discussion and support.

## Supplemental Material

*Acidianus* DS80 was originally isolated from Dragon Spring (78°C, pH 3.1), located in the Hundred Springs Plain area of Norris Geyser Basin in YNP. The initial inoculum for these experiments was obtained from an active pure culture growing anaerobically on H_2_/S^0^ provided by the Boyd lab at Montana State University.

### Preparation of Wolfe’s Vitamins Solution

Wolfe’s vitamins solution was prepared by dissolving the following components in the appropriate volume of ultrahigh purity water: Pyridoxine HCl (0.01g/L), p-aminobenzoic acid (0.005 g/L), lipoic acid (0.005 g/L), nicotinic acid (0.005 g/L), riboflavin (0.005 g/L), thiamine HCl (0.005 g/L), calcium pantothenate (0.005 g/L), biotin (0.002 g/L), folic acid (0.002 g/L), cyanocobalamin (0.0001 g/L). Complete Wolfe’s vitamins solution was filter-sterilized and frozen at −20°C, with one liquid working stock kept in the dark at 4°C.

### Preparation of SL-10 Trace Elements Solution

SL-10 trace elements solution was prepared by dissolving the following components in the appropriate volume of ultrahigh purity water: FeCl_2_ x 4H_2_O (1.5 g/L), ZnCl_2_ (0.07 g/L), MnCl_2_ x 4H_2_O (0.1 g/L), H_3_BO_3_ (0.006 g/L), CoCl_2_ (0.1037 g/L), CuCl_2_ x 2H_2_O (0.002 g/L), NiCl_2_ x 6H_2_O (0.024 g/L), Na_2_MoO_4_ x 2H_2_O (0.036 g/L), Na_2_WO_4_ (0.015 g/L), Na_2_SeO_3_ (0.0098 g/L). The solution was acidified using HCl (33% v/v) at a concentration of 7.5 mL/L. SL-10 trace elements solution was filter-sterilized and stored in the dark at 4 °C.

### Cline assay for tracking S^0^ reduction

The methylene blue method (Fogo and Popowsky, 1949) was used to track sulfide production throughout the course of the experiment. Briefly, 15 μL ferric chloride, 50 μL amine-sulfuric acid, and 750 μL sample (withdrawn from bottles using a 1 mL syringe) were combined in a 1.5 mL centrifuge tube. The solution rapidly turned blue in the presence of dissolved sulfide. Immediately after the color change, 160 μL of diammonium hydrogen phosphate (DHP) was added to quench the reaction, resulting in the formation of a white precipitate. The centrifuge tube was vortexed briefly to dissolve the precipitate. Contents of the centrifuge tube were poured into a 1 mL glass cuvette and absorbance was measured at 670 nm on a Genesys 10S UV-Vis spectrophotometer (Thermo Fisher).

The average dissolved sulfide concentration was calculated from three biological replicates, and the amount of total sulfide produced was calculated assuming standard gas-phase equilibrium. A temperature-corrected Henry’s constant (K_H_) was calculated using a Henry’s constant (K_H_°) of 0.087 and temperature dependence on solubility constant, (–dlnK_H_)/dT, of 2100 (De Bruyn et al., 1995)

### Ferrozine assay for tracking Fe^3+^ reduction

Ferrous iron (Fe^2+^) production was tracked using the ferrozine assay (Viollier et al., 2000). At each timepoint, 300 μL of sample (withdrawn from bottles using a 1 mL syringe) was transferred to a centrifuge tube, and gently spun down at 12xG for 30 seconds to separate out inorganic suspended matter. 100 μL of the resulting supernatant liquid was transferred to a centrifuge tube containing 900 μL ferrozine solution. The mixture turned purple in the presence of Fe^2+^. The tube was vortexed briefly to mix, and contents were poured into a 1 mL glass cuvette. Absorbance was measured at 562 nm on a Genesys 10S UV-Vis spectrophotometer (Thermo Fisher).

---

**Table S1.** Peak areas for GDGT species detected via HPLC-APCI-MS operated in single ion monitoring mode. Ring index is a weighted average of cyclopentane rings in all GDGTs from a sample (see Eqn. 1).

| Sample | GDGT-0 | GDGT-1 | GDGT-2 | GDGT-3 | GDGT-4 | GDGT-5 | GDGT-6 | Avg. Ring Index |
| --- | --- | --- | --- | --- | --- | --- | --- | --- |
| H<sub>2</sub>/S<sup>0</sup>/glucose (A) | 7655 | 21181 | 63525 | 50793 | 81490 | 65327 | 27754 | 3.61 ± 0.11 |
| H<sub>2</sub>/S<sup>0</sup>/glucose (B) | 6957 | 16888 | 50255 | 43737 | 67372 | 74965 | 28543 |  |
| H<sub>2</sub>/S<sup>0</sup>/CO<sub>2</sub> (B1) | 14412 | 14713 | 17423 | 32024 | 111029 | 94416 | 30840 | 3.89 ± 0.12 |
| H<sub>2</sub>/S<sup>0</sup>/CO<sub>2</sub> (B2) | 11199 | 10420 | 26195 | 29942 | 101001 | 95132 | 29950 |  |
| H<sub>2</sub>/S<sup>0</sup>/CO<sub>2</sub> (B3) | 12560 | 11297 | 20975 | 33236 | 70639 | 66302 | 15390 |  |
| H<sub>2</sub>/Fe<sup>3+</sup> (B1) | 8126 | 3006 | 20198 | 46080 | 288336 | 162159 | 40561 | 4.10 ± 0.07 |
| H<sub>2</sub>/Fe<sup>3+</sup> (B2) | 19933 | 13629 | 22052 | 64672 | 406225 | 200677 | 27471 |  |
| H<sub>2</sub>/Fe<sup>3+</sup> (B3) | 19267 | 13218 | 36534 | 56144 | 437411 | 203314 | 44314 |  |
| S<sup>0</sup>/Fe<sup>3+</sup> (B1) | 3600 | 1070 | 5881 | 24615 | 262618 | 288135 | 133555 | 4.71 ± 0.02 |
| S<sup>0</sup>/Fe<sup>3+</sup> (B2) | 861 | 2482 | 3227 | 24833 | 286872 | 328272 | 151953 |  |
| S<sup>0</sup>/Fe<sup>3+</sup> (B3) | 1720 | 1145 | 8121 | 27221 | 268058 | 299149 | 133727 |  |

**Table S2.**
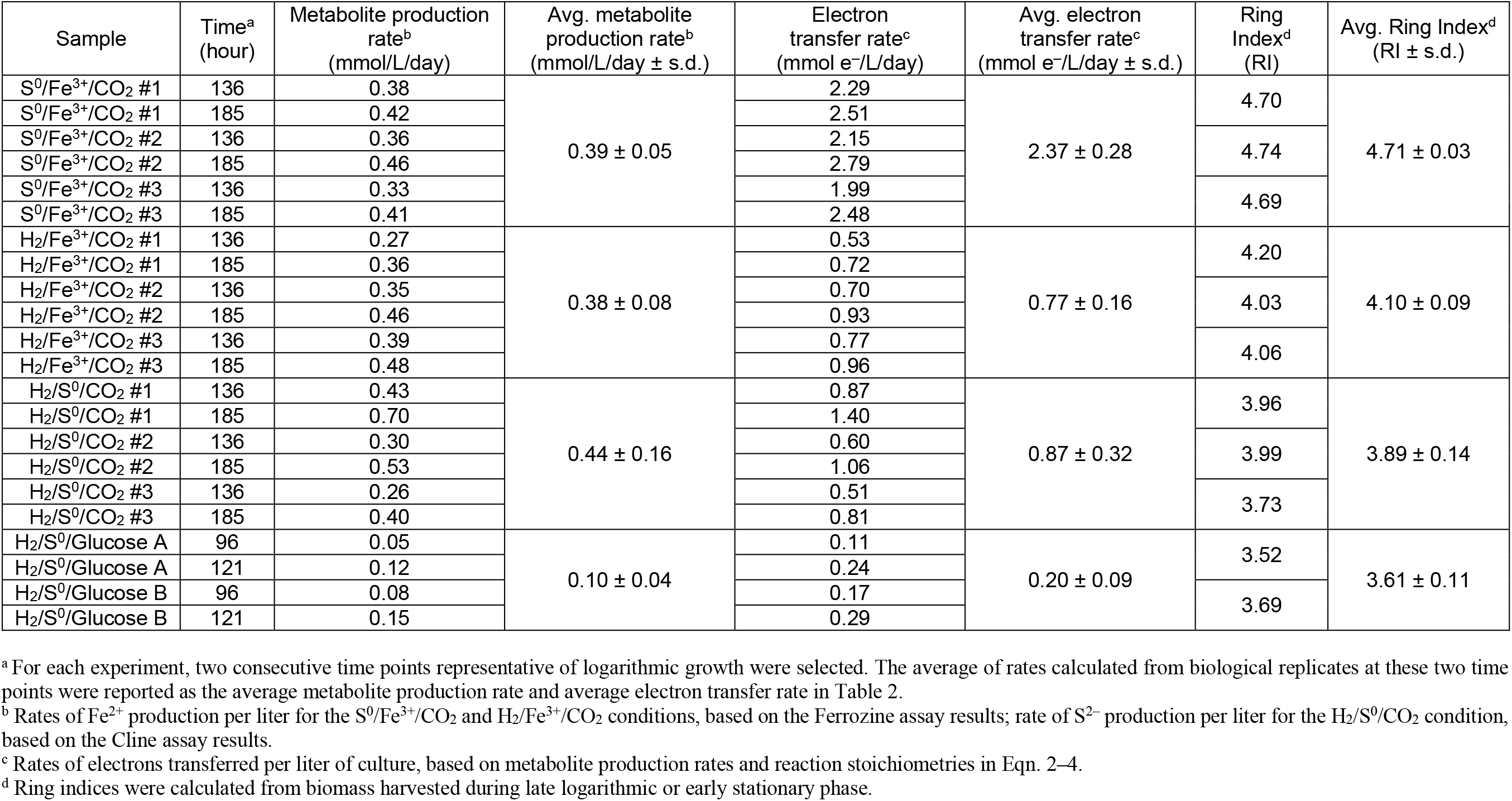
Summary of bioenergetics, metabolic rates, and ring indices across all conditions tested in this study. Rates of metabolites produced and electrons transferred were calculated for data points representing logarithmic growth. Average ring indices were calculated for the biomass harvested during late logarithmic or early stationary phase.

**Figure S1.**
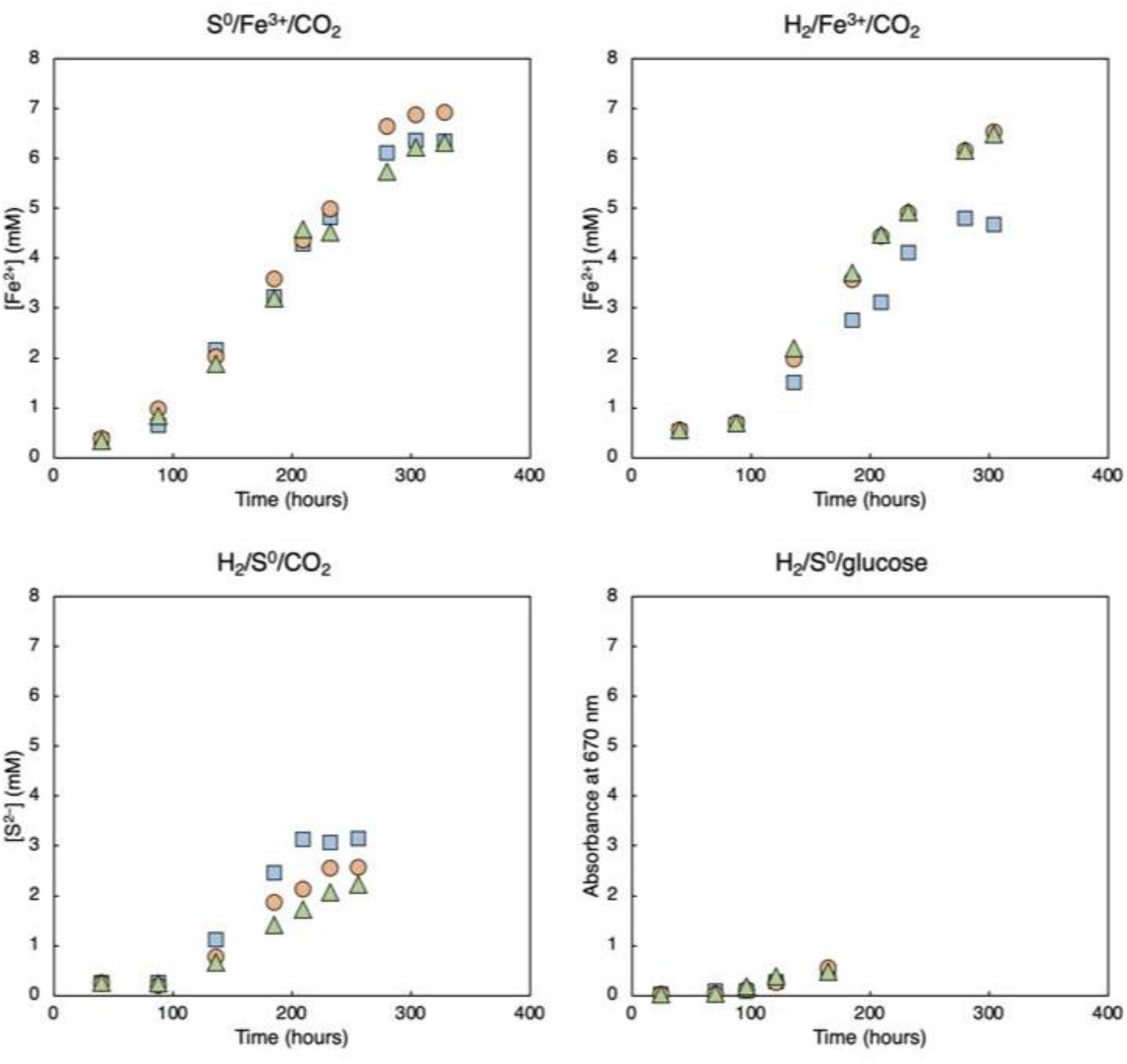
Metabolite production during the growth of DS80. Changes in the concentration of [Fe_2+_] during growth on S^0^/Fe^3+^/CO_2_ (A) and H_2_/Fe^3+^/CO_2_ (B). Changes in the concentration of [S^2–^] during growth on H_2_/S^0^/CO_2_ (C) and H_2_/S^0^/glucose (D). For Fe^3+^-reducing conditions (A and B), the absorbance measured via the ferrozine assay was used to calculate [Fe_2+_]. For S_0_-reducing conditions (C and D), the absorbance measured via the methylene blue assay and standard gas-phase equilibrium calculation were used to estimate total sulfide concentration or [S_2–_]. Biomass was harvested immediately after the final time point, when it was determined that cultures reached an early stationary phase.

**Figure S2.**
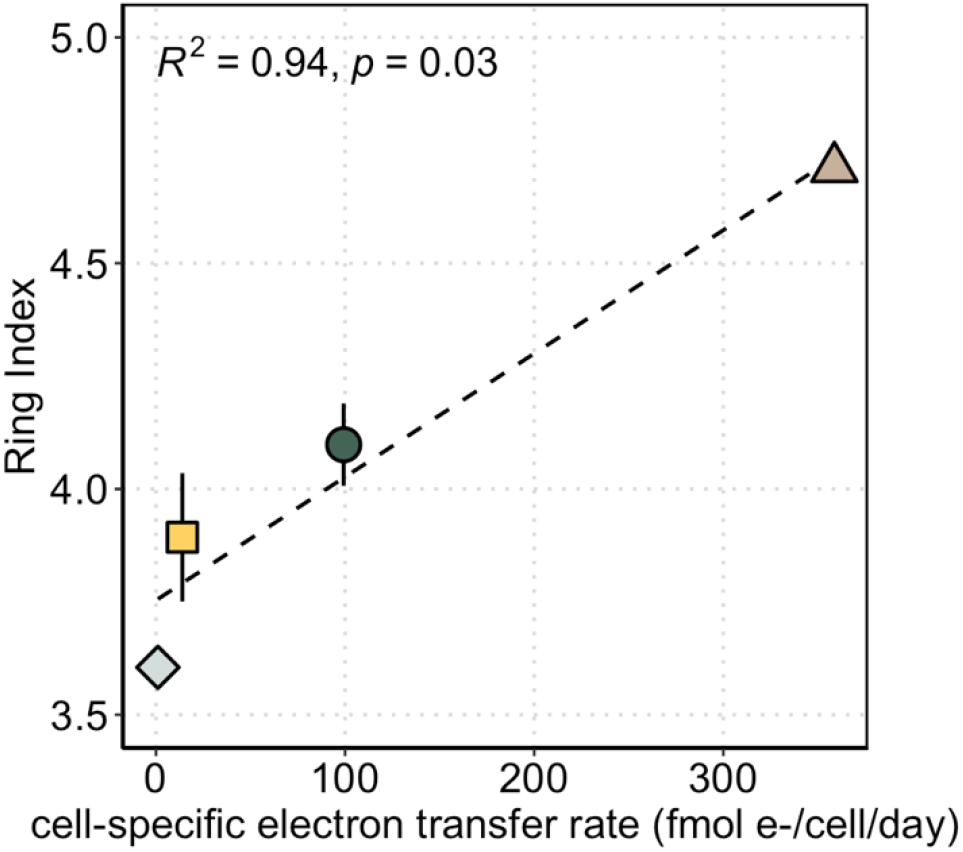
Average ring index values as a function of cell-specific electron transfer rate. The rates were calculated from the measured rates of Fe_2+_ or S_2–_ production per liter of culture, reaction stoichiometries in Eq. 2–4, and the growth yield inferred from the data reported in Amenabar et al. (2017). The same growth yield value for the H_2_/S^0^/CO_2_ treatment in Amenabar et al. (2017) was used for both the H_2_/S^0^/CO_2_ and H_2_/S^0^/glucose treatments in this study.

